# Nucleo-plastidic PAP8/pTAC6 couples chloroplast formation with photomorphogenesis

**DOI:** 10.1101/2020.03.10.985002

**Authors:** Monique Liebers, François-Xavier Gillet, Abir Israel, Kevin Pounot, Louise Chambon, Maha Chieb, Fabien Chevalier, Rémi Ruedas, Adrien Favier, Pierre Gans, Elisabetta Boeri Erba, David Cobessi, Thomas Pfannschmidt, Robert Blanvillain

## Abstract

The initial greening of angiosperm occurs upon light-activation of photoreceptors that trigger photomorphogenesis followed with the development of chloroplasts. In these semi-autonomous organelles, the construction of the photosynthetic apparatus depends on the coordination of nuclear and plastid gene expression. Here we show that PAP8, as an essential subunit of the plastid-encoded RNA polymerase, is under the control of a regulatory element recognized by the photomorphogenic factor HY5. PAP8 is localized and active in both plastids and the nucleus and particularly essential for the formation of late photobodies. In the albino *pap8* mutant, phytochrome-mediated signalling is altered, PIFs are maintained, HY5 is not stabilized, and GLK1 expression is impaired. PAP8 translocates into plastids losing its pre-sequence, interacts with the PEP, and using an unknown route or a retrograde transport, reaches the nucleus where it has the ability to interact with pTAC12/HMR/PAP5. Since PAP8 is required for the phytochrome-B-mediated signalling cascade and the reshaping of the PEP, it may coordinate nuclear gene expression with the PEP-driven chloroplastic gene expression during chloroplast biogenesis.

## Introduction

Chloroplasts are the organelles conducting photosynthesis in plants and green algae (Jarvis & Lopez-Juez, 2013, Liebers, Grubler et al., 2017). In angiosperms, the photosynthetic organelles differentiate from proplastid in a light-dependent manner involving photoreceptors. Seedlings sheltered from light, launch a chloroplast-free developmental programme, named skotomorphogenesis, leading to cell elongation of specific organs (e.g. the hypocotyl). The inhibition of chloroplast development in the dark can be regarded as a way to optimize the use of limited resources stored in the seed to efficiently reach the surface. Then illumination causes the conversion of photoreceptors such as the phytochromes into an active state launching a developmental programme named photomorphogenesis (Solymosi & Schoefs, 2010). This programme involves the repression of hypocotyl elongation and the opening of the cotyledons that are rapidly engaged in chloroplast biogenesis, a tightly regulated process usually regarded as a part of the photomorphogenic programme (Pogson, Ganguly et al., 2015).

As remnant of their endosymbiotic origin, plastids possess their own genetic system, which contributes to the construction of the photosynthetic apparatus after illumination (Jarvis & Lopez-Juez, 2013). A <u>p</u>lastid-<u>e</u>ncoded RNA <u>p</u>olymerase (PEP) is required for proper transcription of photosynthesis genes encoded by the plastid genome. The PEP complex is composed of a prokaryotic core of 4 distinct bacterial-like subunits (α_2_, β, β’, β”) surrounded by (at least) 12 additional nuclear-encoded subunits of eukaryotic origin (Pfannschmidt, Blanvillain et al., 2015) known as <u>P</u>EP-<u>a</u>ssociated <u>p</u>roteins (PAPs). The association of PAPs to the prokaryotic core is strictly light induced and phytochrome-mediated (Pfannschmidt & Link, 1994, Yang, Yoo et al., 2019, Yoo, Pasoreck et al., 2019). Importantly, the PAP association to the core of the PEP appears to be one important bottleneck of chloroplast formation since genetic inactivation of any PAP results in albinism (Pfalz & Pfannschmidt, 2013). The genes for PAPs appear to represent a regulatory unit that exhibits very similar co-expression profiles albeit the involved genes encode proteins that belong to very different functional classes that could not be predictably united before. They all exhibit a basal expression in the dark followed by a rapid and transient peak after light exposure strongly suggesting a connection of their expression to the light regulation network (Liebers, Chevalier et al., 2018).

In the dark, photomorphogenesis is strongly inhibited by the negative regulatory module CONSTITUTIVE PHOTOMORPHOGENIC/DEETIOLATED/FUSCA (COP/DET/FUS) (Sullivan, Shirasu et al., 2003). In particular, the E3 ubiquitin ligase COP1 was shown to destabilize two basic domain/leucine zipper (bZIP) transcription factors (ELONGATED HYPOCOTYL 5, HY5 and its homologous protein HYH) (Holm, Ma et al., 2002, Osterlund, Hardtke et al., 2000). Upon illumination, the cytosolic pool of inactive phytochrome B (PHYB^Pr^) is converted into an active state (PHYB^Pfr^), triggering its translocation into the nucleus (Yamaguchi, Nakamura et al., 1999) where it physically interacts with PHYTOCHROME-INTERACTING FACTORS (PIFs) (Huq, Al-Sady et al., 2004) leading to the emergence of a mutual negative feedback loop (Leivar & Monte, 2014). PIFs, as transcriptional repressors of photomorphogenesis, are destabilized leading to a de-repression of the photomorphogenic programme (Jiao, Lau et al., 2007). In particular, the COP1-mediated destabilization of HY5 is abolished, thereby leading to its accumulation. Stabilized HY5 can then initiate expression of photomorphogenic factors (Lee, He et al., 2007). Meanwhile, light exposure triggers the transcriptional activation of GOLDEN2-LIKE 1 and 2 (GLK1 and 2) transcription factors that are responsible for the proper expression of nuclear photosynthesis genes (Waters & Langdale, 2009, Waters, Wang et al., 2009). The action of phytochrome within the nucleus was visualized using a GFP-tag (PHYB-GFP or PBG) revealing that phytochrome B translocates into the nucleus, and then aggregates into specific speckles within the nuclear matrix (Yamaguchi et al., 1999). Early speckles are numerous and small, while later speckles become larger and less abundant. Late speckle formation, also designated “late photobodies”, requires the presence of HEMERA (HMR), a bi-localized protein present in the nucleus and in plastids (Chen, Galvao et al., 2010, Nevarez, Qiu et al., 2017). In plastids HMR is known as pTAC12/PAP5 representing a member of the PAP family that is essential for chloroplast biogenesis since genetic inactivation of the protein blocks plastid differentiation leading to albinism (Pfalz, Holtzegel et al., 2015, Pfalz, Liere et al., 2006). For PAPs, different functions can be predicted from their amino-acid sequences, but their precise roles are not yet understood. PAP8 is one of the most enigmatic members among the PAPs, as its deduced amino-acid sequence does not harbour any known functional domain. Here we show that PAP8 is a bi-localized nucleo-plastidic protein with a nuclear pool important for the proper timing of chloroplast biogenesis. In particular PAP8 is essential for phytochrome-mediated signal transduction, PIF1 and PIF3 degradation, HY5 stabilization and GLKs transcript accumulation indicating that it represents a novel key regulator of the light-signalling network.

## Results

### PAP8 plays an essential role in chloroplast biogenesis

PAP8 was originally identified by targeted proteomics as pTAC6, a component of the transcriptionally active chromosome of plastids. The T-DNA insertion line ‘SALK_024431’ of *Arabidopsis*, referred as the *pap8-1* allele in this study, displayed an albino phenotype (Fig. S1) with a strong depletion of photosynthesis transcripts and pigments accumulation as well as developmentally arrested plastids (Pfalz et al., 2006). An orthologous protein of pTAC6 was then isolated from a purified *Sinapis alba* PEP complex and subsequently renamed PAP8 as being a *bona fide* component of this PEP complex (Steiner, Schroter et al., 2011). The *pap8-1* allele corresponds to the insertion of an inverted repeat of the T-DNA into the first intron of the gene (Fig. 1A). Amplicon sequencing, after PCR-based genotyping (Fig. 1B), showed that 11 bp of the second exon are missing so that the open reading frame (ORF) is destroyed notwithstanding possible events of T-DNA splicing. Besides, a *PAP8* transcript spanning the insertion point could not be detected with RT-PCR in the homozygote *pap8-1* mutant (Fig. 1C), indicating that *pap8-1* is a genuine null allele. The conceptually translated protein sequence is found in the terrestrial green lineage starting from mosses to Eudicots (Fig. 1D), though absent in ferns, gymnosperms and a few basal angiosperms. A predicted N-terminal chloroplastic transit peptide (cTP) rapidly diverged while a highly conserved region (HCR) of unknown function seems to be under a strong selection pressure, as it is almost unchanged since the last common ancestor of all terrestrial plants. Hence the sporophytic lethality of the *pap8* mutant triggers the assumption that the protein had brought an important function to the green lineage in its way to conquer dry lands, and then became essential to Eudicots.

**Fig. 1.**
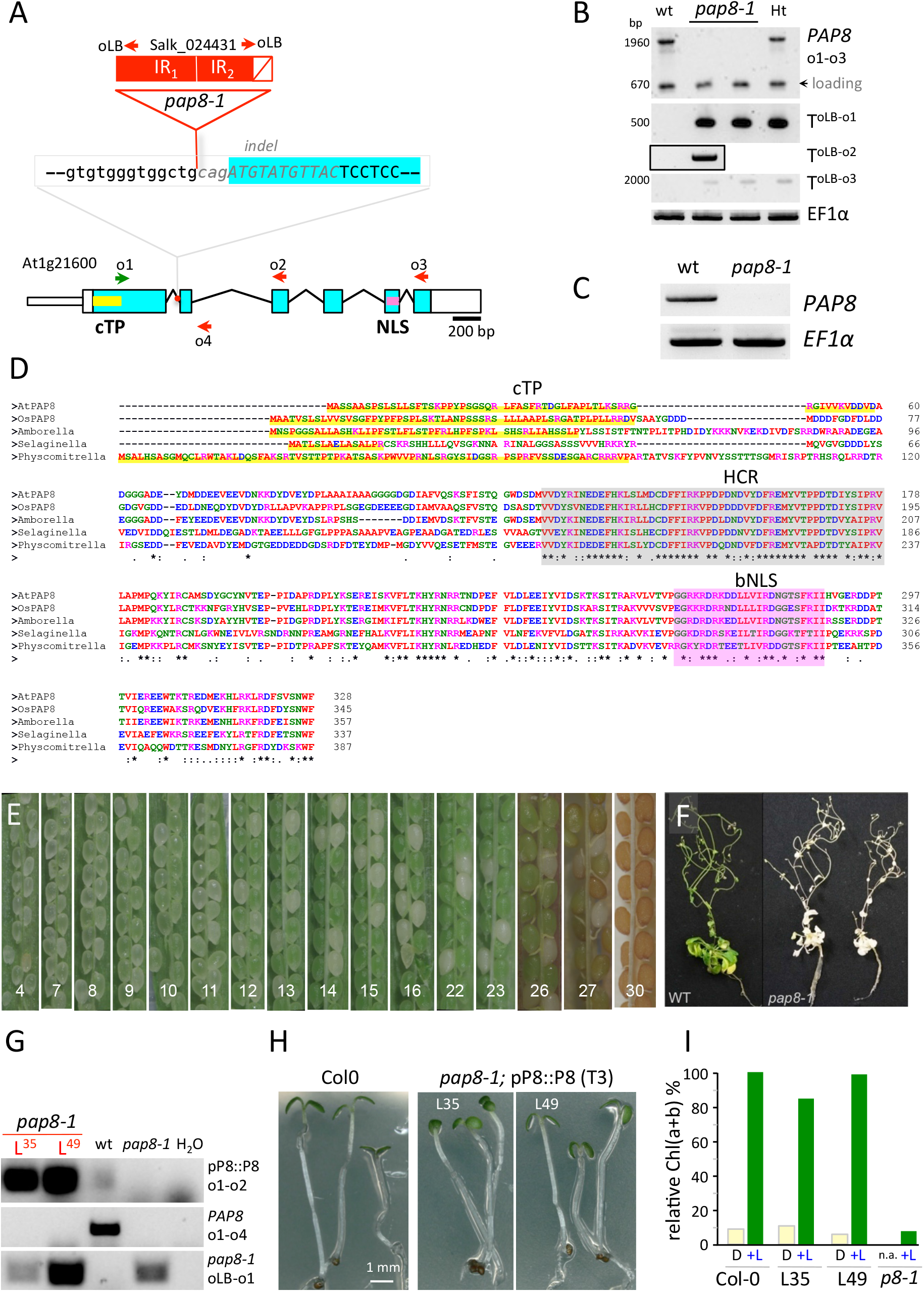
Genetic analysis of the mutant *pap8-1*. **A** Structure of the *PAP8* locus, blue boxes: exons, lines: introns. Red box: inserted T-DNA as inverted repeats (IR_1_/IR_2_) in the first intron. White box with a diagonal red line: deletion at the left border of IR_2_ and part of second exon (italicized grey sequence). **B** PCR performed on genomic DNA with indicated primers as shown: o1: oPAP8_rtpF, o2: oPAP8_E3R, o3: oPAP8_rtpR, oLB: oLBb1.3, WT: wild type, *pap8-1*: Homozygous albino plant, Ht: Heterozygous green plant; T: T-DNA, arrowhead: 670-bp contaminant amplification product used as loading control. **C** RT-PCR on wild type and *pap8-1* homozygous plants grown in the dark for 3 days followed with 72-h growth under white light to allow greening of the wild type; *EF1α* used as control. **D** Sequence alignment of predicted full length orthologous PAP8 protein found in representatives of major phylogenetic clades *Arabidopsis thaliana*, At1g21600; *Oryza sativa Indica*, EEC67529.1; *Amborella trichopoda*, XP_006827378.1; *Selaginella moellendorffii*, XP_002976643.2 *Physcomitrella patens*, XP_024396032.1. cTP, chloroplast transit peptide as predicted with ChloroP1.1 (www.cbs.dtu.dk) underlined in yellow; HCR shaded in grey, highly conserved region. (*), (:), (.), conserved, strongly similar, or weakly similar amino acid properties (standards from www.uniprot.org). bNLS, bipartite NLS as predicted with NLS mapper (http://nls-mapper.iab.keio.ac.jp). **E** Half-open siliques of a heterozygous plant showing the embryo greening. The given number is the position rank of the silique from top to bottom of the inflorescence presenting the segregation of homozygous and heterozygous seeds based on their ability to transiently develop chloroplasts. **F** Pharmacological rescues of *pap8-1 in vitro* using sucrose and low white light intensity of 10 μmol.m^−2^.s^−1^. **G** Representative 8-week-old plants retrieved for the picture. **G-J** Functional complementation of *pap8-1* with pP8:PAP8^cds^: PAP8 coding sequence under control of a 1.1-kb upstream region used as promoter (see Fig. 2a). **G** PCR on genomic DNA; L35, L49: Two independent transgenic lines; primers are the same as in Fig. 1b and o4: op8i2_R. **H** Greening assay on wild type and rescued *pap8-1* homozygous plants grown *in vitro* 3 days in the dark followed with a 30-h light treatment (+L). **I** Content of total chlorophylls (Chl(a+b)) normalized to fresh weight and relative to wild type in the given genotypes grown in the dark (D) or grown in the dark followed with 30 hours of white light treatment (+L); n.a. not applicable.

**Figure 2.**
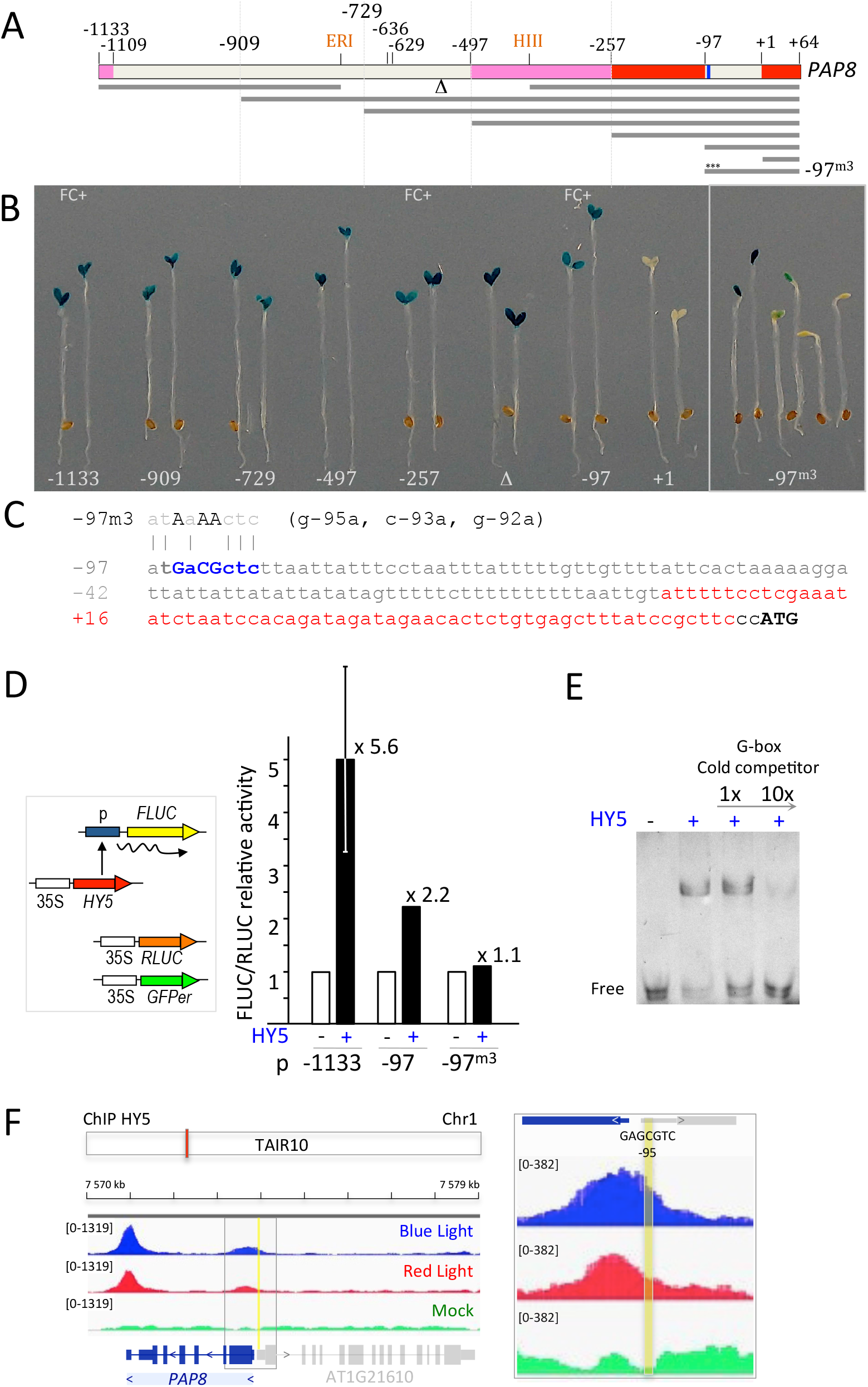
HY5 is a potential regulator of PAP8 expression. **A** *PAP8* promoter deletion strategy. ERI, *Eco*RI site; HIII, *Hind*III site; indicated positions are given relative to the transcription start noted as +1; red boxes represent untranslated regions; pink boxes, ORF of an upstream gene; nearly palindromic element is given in blue. **B** Two representative primary transformants expressing GUS under the given *PAP8* promoter version; FC+, the corresponding promoters were tested positive in functional complementation of the mutant *pap8-1*. **C** Proximal *PAP8* promoter region; m3, 3-bp substitutions within the −97-bp promoter; 5’-UTR in red; ATG, start codon of *PAP8*. **D** Dual luciferase reporter assay; *Renilla* luciferase (Rluc) used as internal control and GFPer used as control for the transfected area; the promoters driving Firefly luciferase (Fluc) were transfected in onion epidermis cells without or with constitutively expressed *HY5*. The Fluc/Rluc activity was set to 1 for the minus-*HY5* control. **E** Electro-mobility shift assay of a probe corresponding to the near palindromic *PAP8* element (GAcGCTC) with recombinant HY5 protein; a probe containing a canonical G-box element (CACGTG) recognized by HY5 was used as cold competitor. **F** Integrative genomics viewer (IGV) images of the Chromatin immunoprecipitation (ChIP) sequencing data^30^ at the PAP8 locus; TAIR10, annotation according to the *Arabidopsis* thaliana information resource. ChIP on *hy5-ks50*; 35S:HY5-YFP exposed to blue light or red light using GFP antibody and compared to mock corresponding to ChIP control experiment done without antibody. Each treatment is presented as track overlay of triplicates: the read count is given within the “group autoscale” range in brackets. Close up on the 5’-UTR region centred on the −95-promoter element in yellow.

In all orthologous proteins a nuclear localization signal (NLS) could be predicted, pointing to a possible localization of the protein inside the nucleus. Functional complementation (Fig. S2) using a full length coding sequence of *PAP8* driven by 1.1 kb of its own promoter (pPAP8∷P8; Table S1) could fully restore the greening of the mutant with a chlorophyll content undistinguishable from that of the wild type (Fig. 1G-I; Fig. S1D-E). Heterozygotes were phenotypically undistinguishable from wild type except within the developing silique where one quarter of the embryos were unable to green (Fig. 1E) following, without gametic distortion, the Mendel’s ratio for the segregation of recessive alleles (Fig. S1B). Mutant homozygotes, however, were albino and sporophytic lethal, with a strong reduction of cotyledon size (Fig. S1C). *pap8-1* dies quickly after light exposure unless grown *in vitro* on a carbon source in dimmed light (Fig. 1F). Albeit their heterotrophic growth, plants pursued a rather normal development until reproduction. PAP8 is therefore a specific factor essential for chloroplast biogenesis without noticeably affecting plastid functions or the apparent photomorphogenic programme that is associated with de-etiolated plants.

### The *pap8-1* promoter involves typical light-responsive *cis*-elements

PAP genes are transcriptionally co-regulated (Liebers et al., 2018, Pfannschmidt et al., 2015); as a canonical example, the promoter activity of *PAP8* is transitorily specific to tissues with photosynthetic potential such as the cotyledons and leaf primordia. It is first restricted to the epidermis during skotomorphogenesis, induced in the palisade after light exposure and then slowly diminished (Liebers et al., 2018). Searching for *cis*-regulatory elements by a deletion series of the *PAP8* promoter, a short sequence starting at −97 from the transcriptional initiation start (tis) was found to be sufficient to retain cotyledon specificity while a construct starting at position +1 completely lost its reporter activity (Fig. 2A, B). The two short versions of the promoter (−257 and −97) driving PAP8 expression were able to complement *pap8-1* (Table S1). Within the 97-bp region (Fig. 2C), a nearly palindromic element (GAcGCTC) was predicted to be a putative non-symmetrical element recognized by proteins with basic leucine zipper domains (bZIP). Site-directed mutagenesis of this element resulted in a disturbed GUS expression (Fig. 2B). Using PlantPAN3 (Chow et al., 2018) three *bona fide* elements for bZIP transcription factors (TF) were predicted in both strands of the DNA (Fig. S3; Table S2). Interestingly the two bZIP TFs, HY5 and HYH are known to be involved in the early steps of photomorphogenesis (Holm et al., 2002, Li, Zheng et al., 2017). Hence a few bZIP TFs, TGA2 as the best prediction according to the two elements found on the plus strand, HY5 and HYH as educated guesses, and bZIP60 as an out-group related to stress response (Iwata, Fedoroff et al., 2008) were tested in a dual luciferase reporter assay (Fig. S4). HY5 proved to be the most efficient, enhancing transcriptional activity of the long (−1133 bp) *PAP8* promoter region by more than 5 fold over the control (Fig. 2D). For the shorter though functional −97-bp-promoter, HY5 promoted transcriptional activity with a 2-fold increase while a 3-bp replacement in the core of the element yielded significantly reduced activation. Moreover recombinant HY5 was able to specifically bind the *cis*-regulatory element *in vitro* (Fig. 2E) in strength comparable to that of the canonical G-box element used as competitor (Yoon, Shin et al., 2006). In addition, the release of Chromatin-Immuno-Precipitation (ChIP) sequencing data using “GFP” antibody on a *hy5*/HY5∷HY5-YFP genetic background (Hajdu, Dobos et al., 2018) allowed the detection of HY5 on the 5’-region containing the identified regulatory element and the 3’-region of PAP8 after blue light or red light exposure (Fig. 2F). While the expression of *PAPs* is essential for greening, *hy5* mutants display slight greening defects indicating that functional redundancies and compensations occur in the regulation of its target genes (Gangappa & Botto, 2016). For example, the paralogous transcription factor HYH (Holm et al., 2002) is also active on the *PAP8* promoter (Fig. S4). In conclusion ChIP and EMSA indicate that HY5 can bind the PAP8 promoter and that it can activate the promoter in a heterologous system, but given that no expression changes were seen in a *hy5-1* mutant, possibly due to functional redundancy, the ChIPseq/EMSA/transactivation data remains to be challenged in more sophisticated genetic backgrounds. Moreover, the epidermal specificity of the PAP-promoter activity during skotomorphogenesis may result from a separate pathway linked to cell identity in relation to development. In this context though, it is of interest to note that *PHYB* promoter activity in the dark shows a pattern similar to that of the PAP8 promoter (Somers & Quail, 1995).

### PAP8 functions in plastids and in the nucleus

PAP8 displays a predicted chloroplastic transit peptide (cTP) and a predicted nuclear localization signal (NLS) (Pfannschmidt et al., 2015), and therefore may belong to a group of bi-localized proteins (Krause, Oetke et al., 2012). Both signals are simultaneously functional since a translational fusion of PAP8-GFP (Fig. 3A) displayed a bi-localized pattern in nucleus and plastids of transiently transfected onion cells (Fig. 3B). A polyclonal serum was raised against a recombinant protein PAP8 conceptually processed, starting after the predicted cleavage site. The specificity of the serum was validated *in planta* using the mutant *pap8-1* as well as the recombinant protein (rP8, Fig. S5A). PAP8 is largely enriched in the sub-cellular fraction corresponding to sedimented organelles (mostly nuclei and plastids) obtained from 5-day-old *Arabidopsis* seedlings (Fig. S5B). PAP8 was then detected both in the nucleus and the plastid fractions obtained from seedlings either grown in the dark or under white light (Fig. 3C) confirming the dual localization of PAP8. The distribution of PAP8 between the nucleus and the corresponding plastid-type fraction is changed after light exposure. In etiolated seedlings PAP8 was found mainly in the nucleus with traces in etioplasts (EP) while in photomorphogenic seedlings PAP8 was strongly enriched in chloroplasts (CP). Notably, both fractions (nuclei and plastids) displayed a signal of the same apparent molecular weight and similar to that of the designed ΔcTP recombinant protein suggesting that the nuclear fraction contains the processed version of the protein originating from plastids where the cleavage of the pre-sequence occurs during import.

**Figure 3.**
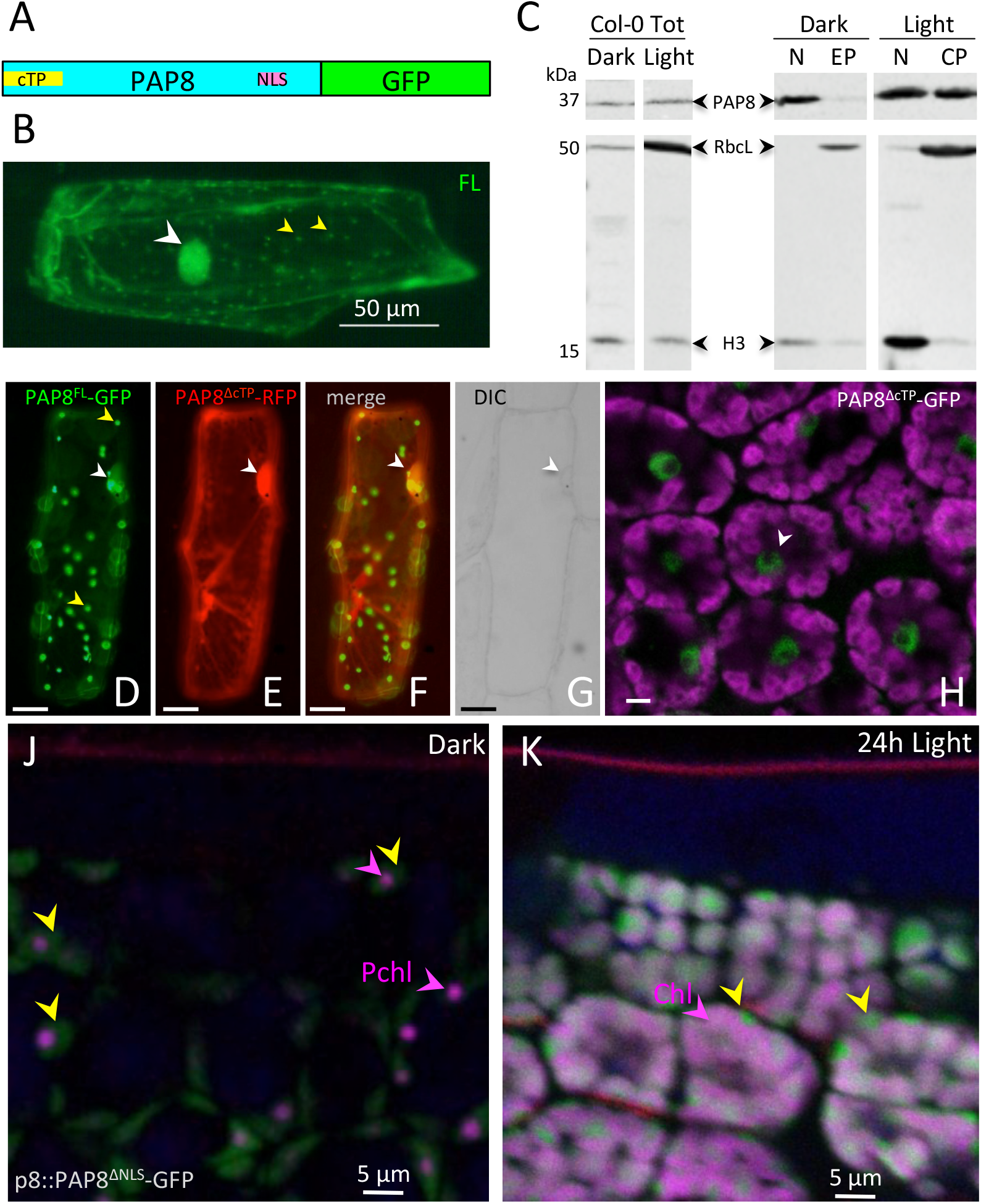
PAP8 is dually localized in plastids and nucleus. **A** Schematic illustration of domain structure of *Arabidopsis* PAP8 fused to GFP. cTP, chloroplast transit peptide; NLS, nuclear localization signal. **B** Transiently expressed PAP8^FL^-GFP (full-length coding sequence of PAP8 fused to GFP) in onion epidermal cells displays a dual localization in the nucleus (white arrowhead) and in plastids (yellow arrowheads). **C** Immuno-blots for the detection of PAP8 in total protein extracts (Col-0 Tot) the nuclear fraction (N) and the plastidic fractions (EP, etioplast; CP, chloroplast) of etiolated (Dark) or photomorphogenic (Light) *Arabidopsis* seedlings by immuno-western blotting using a PAP8 antiserum (PAP8). As control, a mixture of antisera raised against Histone 3 (H3) and the large subunit of ribulose 1, 5-bisphosphate carboxylase/oxygenase (RbcL) was used to evaluate reciprocal contaminations. The lanes are extracted from the same blots provided in the Source Data file. **D-G** Co-expression analysis of PAP8^FL^-GFP **(D)** with PAP8^ΔcTP^-RFP **(E)**, **F** green and red channels merged; white arrowheads show nuclei as observed with DIC, differential interference contrast in **(G)**; FL, PAP8 full length ORF; ΔcTP, deletion of the cTP; bars equal 20 μm. **H** Confocal imaging on *Arabidopsis* cotyledons expressing PAP8^ΔcTP^-GFP, details of a few palisade cells (see Fig. S6B for a view in a cross section between the two cotyledons); magenta, auto-fluorescence of the chloroplast; scale bar equals 5 μm. **J-K** Confocal imaging on *Arabidopsis* cotyledons stably expressing pP8∷PAP8^ΔNLS^-GFP; ΔNLS, deletion of the NLS **(J)** during skotomorphogenesis and **(K)** after 24h light; Yellow arrowheads show the GFP signal; the picture is a merge of different channels: GFP in green, Pch,: protochlorophyllide, or Chl: chlorophyll in magenta marked with arrowheads, and propidium iodide, showing the waxy cuticle in red, the empty space correspond to the layer of highly vacuolated epidermal cells.

To investigate this in more detail PAP8 localization was artificially uncoupled using a mutation strategy. Variants of PAP8-GFP lacking the cTP (ΔcTP), the NLS (ΔNLS), both signals (ΔΔ), or containing a mutated NLS with five neutral substitutions of the positively charged amino acids within the NLS (NLS^m5^) were cloned. In transiently transfected onion cells as well as in *Arabidopsis thaliana* lines with stable expression, PAP8^ΔcTP^-GFP displayed nuclear accumulation (Fig. 3E, H, S6A, B) whereas the ΔNLS and the NLS^m5^ variants were strictly restricted to plastids (Fig. 3J-K, 4A, S6A, C) indicating that the cTP supports chloroplast import and that nuclear localization depends on its NLS. Thus the PAP8 sub-cellular localization can be controlled in transgenic plants using the different targeting signals, and the corresponding transgene can therefore be assessed for functionality in *pap8-1*. Hence PAP8 variants fused or not to GFP, as indicated, were expressed under the constitutive promoter CaMV35S or its own promoter pP8 (pPAP8^−1133^), in wild type or in *pap8-1*. In contrast to pP8∷PAP8, all genetic constructions with the GFP tag were unable to yield functional complementation (Fig. S7). Since GFP may very likely impose a steric hindrance to the function of PAP8, only protein accumulation and subcellular localization were tested using the fluorescent marker. In addition, the greening of plants expressing part or full-length sequence of PAP8 under 35S promoter was strongly altered with no regard to its functionality or its proper localization (Fig. S7b), suggesting that over-expression or miss-expression of the transgene with part of the PAP8 sequence might titter a component of unknown nature (protein, RNA, or else) that affects the biogenesis or stability of the chloroplast within the cell.

**Figure 4.**
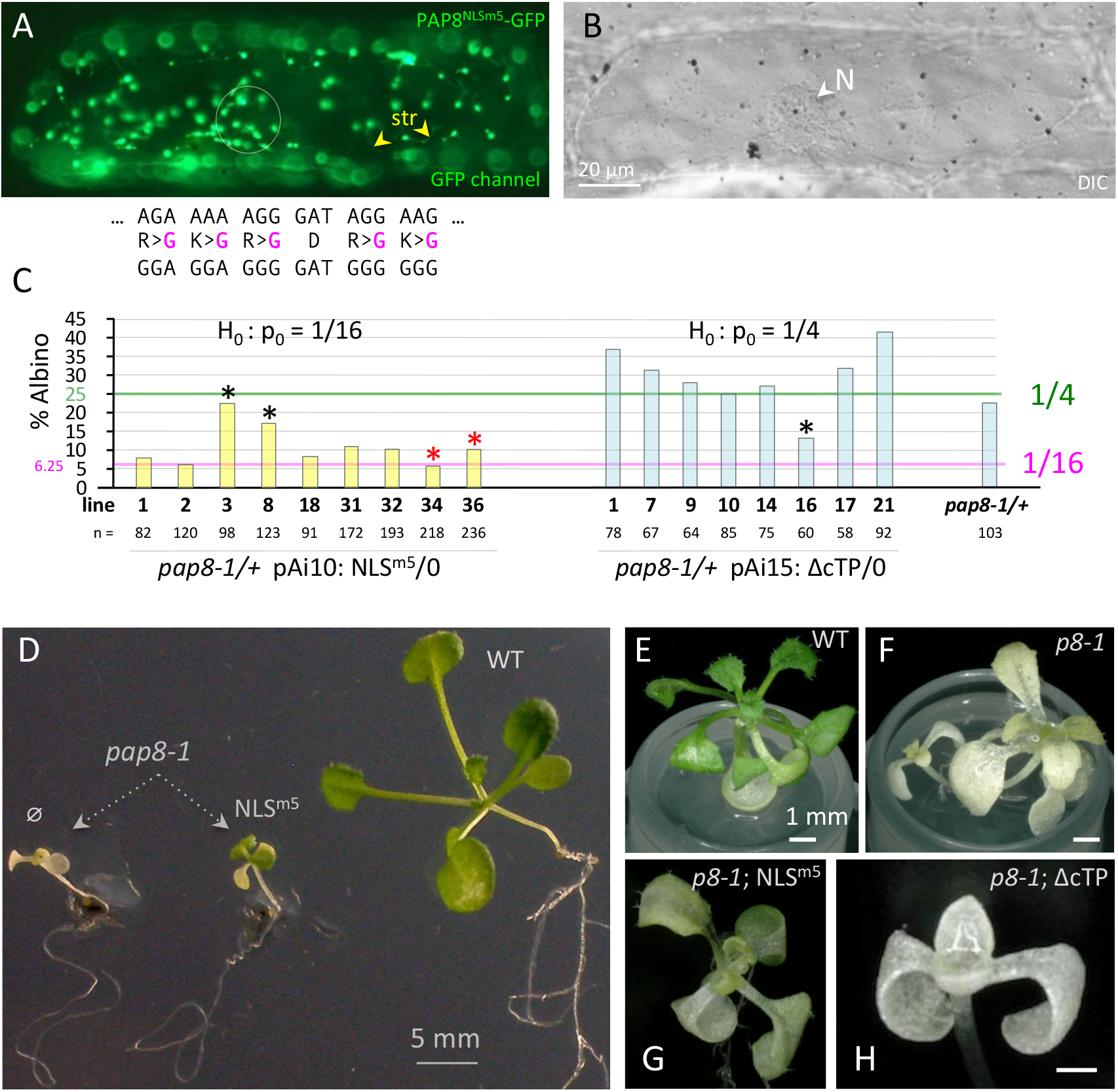
PAP8 is functional in both the chloroplast and the nucleus. **A** Transient expression of PAP8^NLSm5^-GFP mutated in the NLS as noted above. The circle marks the position of the nucleus as observed with DIC in **B** Scale bars equal 20 μm. str, stromules; N, nucleus. **C** Bar graph representing the albino segregation ratios in doubly heterozygous transgenic lines (pap8-1/+; TG/−) obtained with pPAP8∷PAP8NLSm5 (NLSm5: pAi10) and pPAP8∷PAP8ΔcTP (ΔcTP: pAi15). *pap8-1*/+ used as control; n, total number of recorded plants. Segregation pattern were tested using ɛ-test with null hypothesis set to p0= 1/16 (greening complementation) for NLSm5 and p0=1/4 (no complementation) for ΔcTP (see data source); *, outliers correspond to the samples that did not pass the statistical test. **D-H** Pictures of representative genotypes obtained in the functional complementation test of *pap8-1* using pP8∷PAP8-NLS^m5^ or pP8∷PAP8^ΔcTP^ without GFP tags. Pictures using Keyence technology of plants with genotypes as labelled; transgenes expressed using the 1.1-kb PAP8 promoter.

In wild type, the GFP signal of pP8∷PAP8^ΔNLS^-GFP increased in chloroplast sub-domains during the transition from dark to light (Fig. 3J, K). Therefore protein accumulation follows the promoter PAP8 induction in the palisade cells (Liebers et al., 2018), and is consistent with the immune-detection of the native PAP8 in subcellular fractions. In contrast to the fluorescently tagged PAP10 that does not contain a predicted NLS and show a sharp and distinct localization (Fig. S6D), a wider signal of PAP8-GFP indicates that foci slowly appear after light exposure while part of the pool remains in the stroma. The foci, specifically marked with PAP10, may correspond to the assembly of the prokaryotic PEP core complexes with the eukaryotic PAPs. The stroma-localization of PAP8-GFP was confirmed by the PAP8-GFP signal transiently observed in stromules of onion cells while the PAP10-RFP signal is absent from these stromules (Fig. S6E-H). Therefore, PAP8 may be set free from the PEP-PAP complex allowing for re-localization in the nucleus. Whether this release is allowed through saturation of the complex or a change in its affinity remains unknown.

Since GFP-tagged PAP8 could not rescue the mutant, untagged PAP8 variants were tested in functional hemi-complementation (Fig. 4C-H). In contrast to the ΔcTP variant unable to cross the plastidial envelope and unable to rescue the albinism (Fig. 4C, H), the NLSm5 variant could restore the greening of the mutant albeit with strong delays in growth (Fig. 4C-G, Fig. S8) suggesting that the chloroplast-localized PAP8^NLSm5^ carries its chloroplast function for the greening but that in absence of the nucleus-localized pool the timing of chloroplast biogenesis is altered, with substantial consequences on the timing of light-controlled development. Therefore, PAP8, through its nuclear pool, may carry a function related to the light signalling response.

### PAP8 mediates phytochrome signalling

To test this assumption, different light qualities were applied to the plants. Although, in our *in vitro* growing conditions, *pap8-1* responded normally to red and white lights with proper de-etiolation with cotyledon and apical hook opening, far-red light treatment yielded a significantly reduced repression of hypocotyl length, a phenotype similar to that of *hmr-2/pap5-2* (Chen et al., 2010) (Fig. S9).

Stable over-expression of a phytochrome PHYB-GFP (PBG) is known to mediate hypersensitivity of *Arabidopsis* seedlings to red light (8 to 30 μmol.m^−2^.s^−1^, Fig. 5A, B) leading to a significant inhibition of hypocotyl elongation when compared to WT (Yamaguchi et al., 1999). After introducing PBG into the *pap8-1* mutant background, however, this PBG effect was largely lost, indicating that PAP8 plays a role in the PHYB-mediated light response. This lack of physiological response correlates with the retention of small PBG speckles in *pap8-1* corresponding to the absence of late photobodies in comparison to wild type (Fig. 5C-G). The change in the photobodies patterning is not due to a change in PBG accumulation as tested by immune-detection of the GFP tag in the different genetic backgrounds (Fig. 5H). Late photobodies are known to be associated with the targeted degradation of PIFs (Leivar & Monte, 2014), the key regulators in the phytochrome signalling network. This pointed to an active role of PAP8 in the light-induced gene expression programme as illustrated with the strong defects in the accumulation of *GLK1, and GLK2* transcripts in *pap8-1* (Fig. 5I) thus interrupting the light-induced expression of *PhANGs* (Oh & Montgomery, 2014, Waters & Langdale, 2009). The defect in GLKs transcript accumulation was also observed in *pap7-1* (Grubler, Merendino et al., 2017) accounting for the albino syndrome of the *pap* mutants where the expression of *PhANGs* is strongly altered.

**Figure 5:**
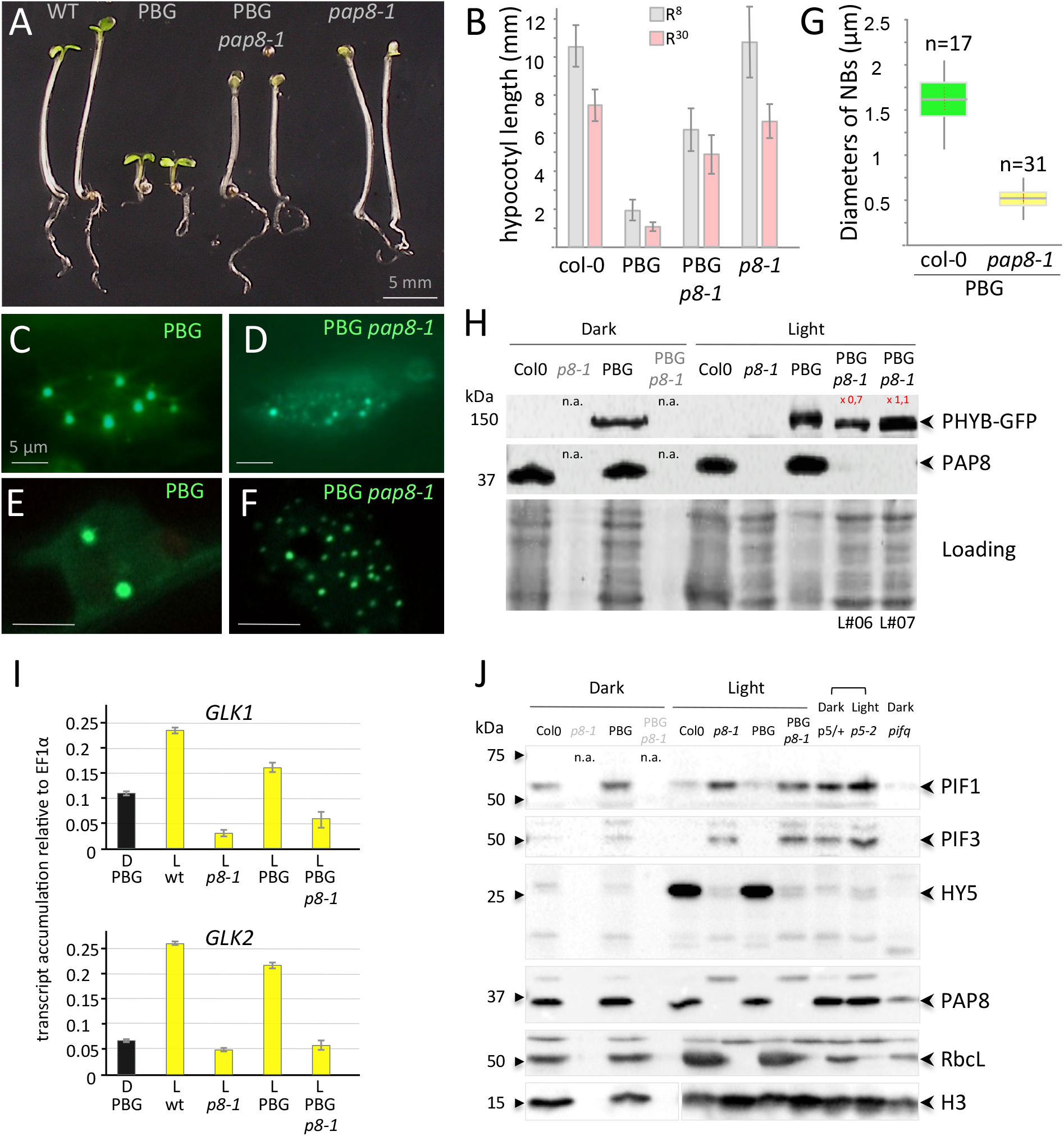
PAP8 is essential for the PHYB-mediated light induction of photomorphogenesis. **A** Phenotypes of given genotypes subjected to 5 days of illumination at 8 μmol.m^−2^.s^−1^ 660-nm red light. PBG, pCaMV35S∷PHYB-GFP transformed in *pap8-1/+* (among 25 lines selected for GFP expression see data source; 2 doubly heterozygous *pap8-1*/+; PBG/− lines #6 and #7 segregated the photobodies alteration with the albinism). **B** Hypocotyl length of plants grown as in **A** (R8, grey bars) or at 30 μmol.m^−2^.s^−1^ 660-nm red light (R30, pink bars) showing partial insensitivity of *pap8-1* to the PBG overexpression (R8: δ^PBG/PBGp8-1^= 20.38>> 1.96 > δ^wt/p8-1^= 0.5). **C-G** Nuclear accumulation of PBG observed under GFP excitation in the given genotypes. **C-D** Epi-fluorescence microscopy. **E-F** Confocal microscopy showing the size of the nuclear bodies. **G** Box plot on the diameter of the nuclear bodies (NBs); n equals the number of records. **H** Immuno-blots using a GFP antibody or a PAP8 antibody showing respectively the levels of PHYB-GFP and PAP8 in the given genotypes grown in the dark for 3 days or in light; n.a., not applicable as the *pap8-1* mutant can only be visually distinguished from wild type after light exposure; 2 lines (L#06 and L#07) for PBG/*pap8-1* were tested. Coomassie blue staining presented as loading; signals were quantified using ImageJ. **I** RT-qPCR analysis on wild type, *pap8-1*, PBG and PBG *pap8-1*. Seedlings were grown in the dark (D) or under white light (L, 30 μmol.m^−2^.s^−1^); levels of transcripts are given relative to EF1α; error bars correspond to standard errors on technical triplicates and the dark sample is the wild-type PBG line. **J** Immuno-blots showing the levels of PIF1, PIF3, HY5 in given genotypes: *p5/+*, mix of an heterozygous *pap5-2* siblings progeny undistinguishable from wild type; p5-2, *pap5-2* and *pifq*, quadruple *pif1-1 pif3-3 pif4-2 pif5-3* mutant; Histone H3 (H3) RbcL and PAP8 were used as controls; n.a., not applicable.

The light-induced destabilization of PIF1 and PIF3 is altered in *pap8-1* and PBG/*pap8-1*, conversely the light-induced stabilization of HY5 does not occur in *pap8-1* (Fig. 5J). Interestingly, these molecular phenotypes in *pap8-1* are very similar to those observed in *pap5-2* used as control. Therefore, PAP8 supports the degradation of PIF1 and PIF3 and stabilizes HY5 with no effect caused by the presence of PBG. In the light signalling cascade PIFs are known to act upstream of HY5 and in a reciprocal negative feedback loop with PHYB (Leivar & Monte, 2014). This indicates that the alteration of the signalling in *pap8-1* (the albino block depicted in Fig. 7) acts upstream of PIF1 and PIF3 by specifically blocking the HY5 to GLK pathway without altering the de-etiolating pathway: the apical hook and the cotyledons can open. The nature of the block remains unknown; it could be due to direct functional alteration of the PAP nuclear sub-complex in which PAP8 and PAP5 may act co-ordinately and dependently, or due to an upstream retrograde signal coming from the challenged *pap8*-deficient chloroplast (Martin, Leivar et al., 2016). Concerning the growth of the hypocotyl, the situation remains complex whether PBG is considered or not.

Therefore, PAP8 is important for the proper expression of *GLKs*. Should this occur through the nuclear function of PAP8, directly or through a PAP8-containing complex, this would simply explain the delayed greening and growth observed in the partially rescued phenotype of the PAP8^NLSm5^ variant, in which nuclear PAP8 is absent. Should the expression of GLK1 be controlled by the state of the plastids through a distinct molecular pathway, this would then be an indirect consequence of the *pap8-1* phenotype and more generally of the *pap* albino syndrome. Future research will probably help solving this conundrum.

### PAP8 physically interacts with HMR/PAP5

The cellular distribution of PBG and other defects in *pap8-1*, are highly similar to those of *hmr-2/pap5-2*. HMR/PAP5 is a nucleo-plastidic protein identified to be (1) important for the initiation of photomorphogenesis (Chen et al., 2010, Qiu, Li et al., 2015), and (2) a component of the chloroplast PEP complex (Nevarez et al., 2017, Steiner et al., 2011). Although yeast-two-hybrid studies did not report any interaction between the two proteins (Arsova, Hoja et al., 2010, Gao, Yu et al., 2011, Yu, Lu et al., 2013), bimolecular fluorescence complementation technology (BiFC, Fig. 6A, S10 for control experiments) revealed that PAP8^ΔcTP^ and HMR/PAP5^ΔcTP^ could together restore split YFP fluorescence indicating that they get in close proximity within the nucleoplasm.

**Fig. 6.**
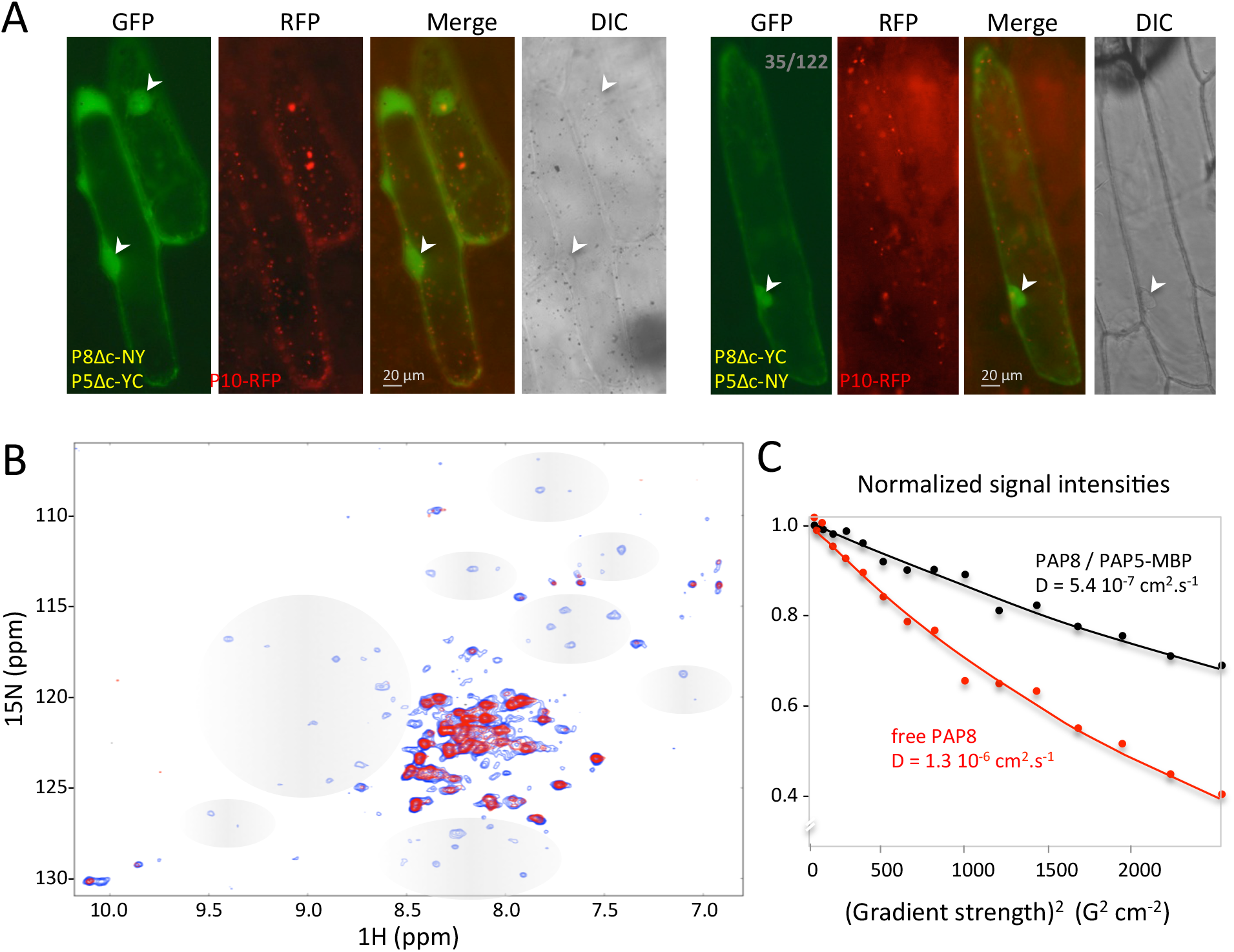
PAP8 interacts with pTAC12/HMR/PAP5. **A** Bimolecular fluorescence complementation tests using in combination PAP8^ΔcTP^-NY (P8Δc-NY) with PAP5^ΔcTP^-YC (P5Δc-YC) or PAP8^ΔcTP^-YC (P8Δc-YC) with PAP5^ΔcTP^-NY (P5Δc-NY); PAP10-RFP (P10-RFP) was used as internal positive control for transfection; arrowheads indicate nuclei. See Fig. S12 for control experiments; transgenes expressed under CaMV35S promoter. **B** Overlay of ^1^H-^15^N correlation 2D NMR spectra of free ^15^N-labelled PAP8 alone (blue) or in complex with PAP5 (red). Grey areas depict changes of signals in the PAP8 spectrum. **C** 15N-Filtered Diffusion Ordered Spectroscopy-NMR measurements to PAP8. Exponential decay curves of PAP8 in absence or in presence of MBP-PAP5 are shown in red and black respectively. The units on the y-axis are normalized values of the integrals of the signal measured in the amide proton region.

The ^1^H-^15^N-correlation NMR spectrum of PAP8 showed two populations of peaks according to their intensities and frequency distributions (blue signal, Fig. 6B). About 40 peaks of high intensities present in a narrow frequency range in the proton dimension (8.0 to 8.5 ppm) correspond to very dynamic and flexible regions of the protein, with a short apparent rotational correlation time (τ_c_ = 2.5 ns, measured at 300 K in a [^15^N,^1^H]-TRACT experiment). The other population of lower intensity peaks with a large frequency distribution corresponds to well-folded domains (τ_c_ = 17 ns). The interaction with PAP5 was tested in a second ^1^H-^15^N-correlation NMR spectrum with unlabelled PAP5-MBP (red signal). The flexible regions of PAP8 were not affected, neither in chemical shift nor in dynamics (τ_c_ ≃ 2.5 ns), and only weakly in intensity. Hence these flexible regions are not involved in the interaction. By contrast, the low intensity peaks from the structured region did not appear when PAP5-MBP was added even after a 14-fold longer experiment (13 h versus 53 min for the free ^15^N-PAP8 spectrum) indicating that PAP5-MPB interacts with the structured region of PAP8.

The robust NMR signals of the PAP8 flexible residues permitted translational diffusion measurements in ^15^N-filtered DOSY spectra with a selective detection of the amide protons bound to the PAP8-^15^N atoms avoiding perturbations by the unlabelled protein. The diffusion rate of free PAP8 was significantly larger than that of the mixture with PAP5-MBP, indicating again that both proteins interact with each other (Fig. 6C). The control experiment using MBP yielded super-imposable spectra and identical PAP8 translational diffusion, indicating that MBP is not involved in the interaction between PAP8 and PAP5-MPB. The Kd corresponding to the interaction between PAP8 and PAP5 was estimated in the range of 50-to-100-μM according to a sub-stoichiometric titration experiment (Fig. S11) (Williamson, 2013).

HMR/PAP5 physically interacts with PAP8 through a well-structured region. Hence, these physical properties reinforce the assumption that PAP8 and PAP5 might form a nuclear complex. It is very likely that additional components could stabilize the unstructured region of PAP8 and enhance its affinity to PAP5 allowing BiFC detection *in vivo*. Should such a nuclear complex exist, the proteins could work cooperatively in an interdependent fashion. Consequently, individual mutant phenotypes would resemble each other as it is observed for *pap8-1* and *pap5-2* reminding genetic epistasis where the lack of one gene is equivalent to the lack of the second gene.

### Concluding remarks

This study revealed that PAP8 represents a novel regulatory component that links photomorphogenesis and chloroplast biogenesis through its dual localization. PAP8, therefore, is a novel member of the nucleo-plastidic protein family involved in chloroplast biogenesis (Yang et al., 2019, Yoo et al., 2019). It is proposed that the nuclear fraction of PAP8 is essential to properly transduce the light signal from photo-activated PHYB to the expression of *GLK1*, one of the master regulators of nuclear photosynthesis genes. The amount of immunodetected PAP8 and PAP5 in dark-grown seedlings rises to nearly their maximum amount within 5 minutes following light exposure (Fig. 7A, B) while the stabilization of HY5 takes a few hours to be detected, closely followed with the rise of the ribulose-1,5-bisphosphate carboxylase/oxygenase. Taken together, the results presented in this study prompted a model of PAP8 action within the transition from skotomorphogenesis to photomorphogenesis (Fig. 7C). A dark operating unknown transcription factor (TF?) allows the production of PAP8 in the epidermal cells where it mostly accumulates in the nucleus (Fig. 3C) in its processed form, suggesting that it passes through the plastid for removal of the transit sequence. Nuclear PAP8 may interact with HMR/PAP5 in a PAP nuclear sub-complex PAP-NSC. Upon light exposure, rapid photo-converted PHYB requires the PAP-NSC to transduce the signal to PIF1 and PIF3 for their COP1-mediated degradation. At this early stage, a non cell-autonomous signal such as one that operates for de-etiolating hypocotyls may then allow the signal to invade the palisade tissue where PIFs are destabilized and HY5 is stabilized escaping COP1-mediated degradation. In turn, HY5 may activate the PAP8 promoter (and potentially other PAP promoters) in cells with a fate associated with photosynthesis, allowing for the assembly of the PEP-PAP complex, itself necessary for the expression of the photosynthesis associated plastid genes (PhAPGs). HY5 was also found on the chromatin associated with both GLK1 and GLK2 (Hajdu et al., 2018) and could therefore activate them directly or indirectly through other light responsive factors. In turn GLKs, under GUN1-mediated retrograde signalling (Tokumaru, Adachi et al., 2017), activate the photosynthesis associated nuclear genes (PhANGs). Both PhANGs and PhAPGs participate safely in the build-up of the photosynthetic apparatus (PS). In the absence of PAP8, PIFs are less degraded, HY5 is not stabilized and the GLK pathway does not operate. Concomitantly PAP8, marked absent, cannot assemble in the PEP-PAP complex and PEP-dependent genes are not correctly expressed. In consequence, chloroplasts do not differentiate leading to the albino syndrome. Whether *PAP8* is a positive signal of retrograde signalling remains to be investigated.

**Fig. 7:**
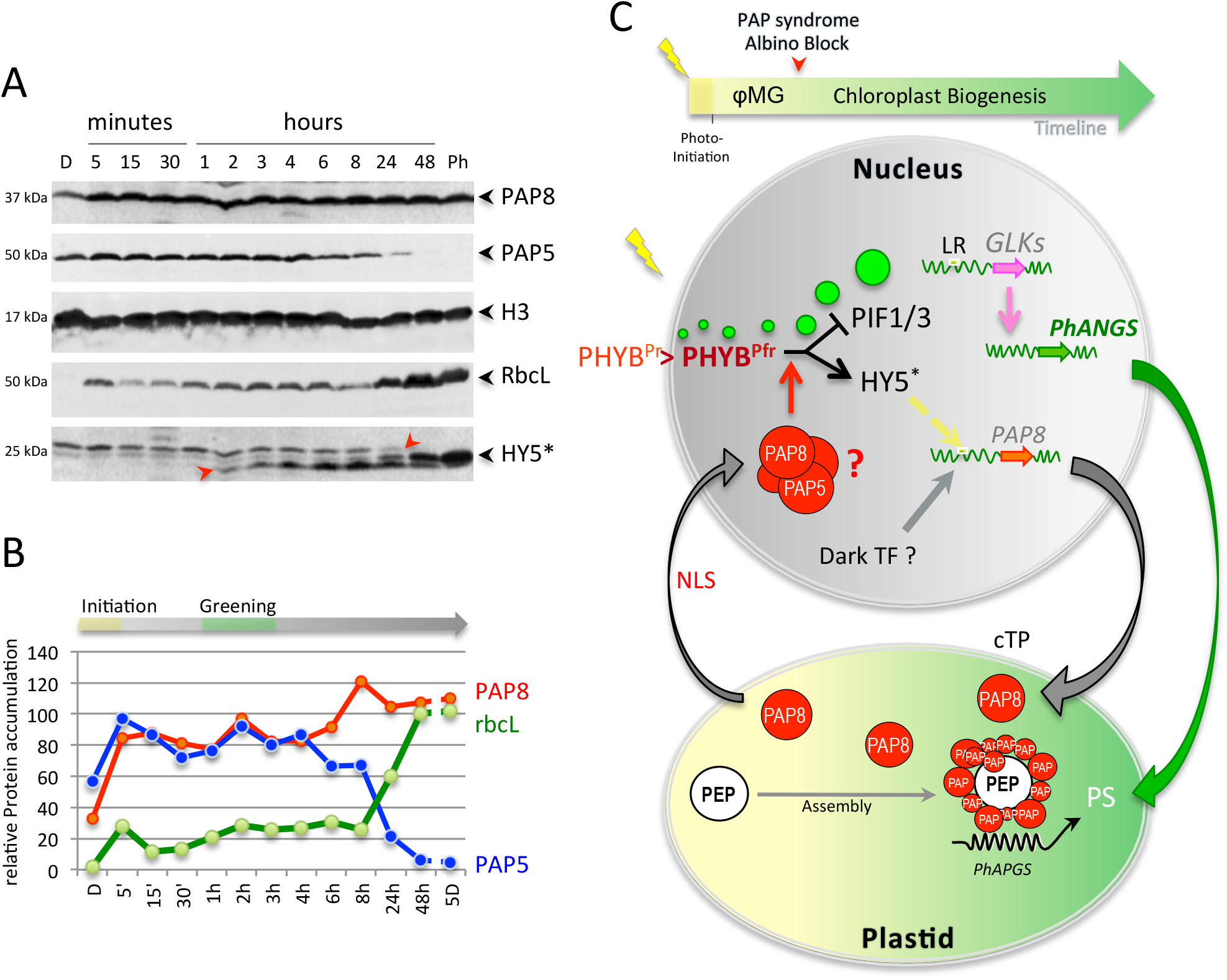
Temporal resolution of protein content in seedlings during the dark-to-light transition. **A** Immuno-blots showing the levels of PAP8, PAP5, Histone H3 (H3), and HY5; D, Dark; Ph, photomorphogenic growth conditions from germination on. **B** Relative protein contents normalized to histone H3 during the transition from skotomorphogenesis to photomorphogenesis, we propose a phase of photo-initiation corresponding to the 5 minutes of light allowing PAP8 and PAP5 to rise to nearly 100% of their maximum. The first macroscopic signs of greening are indicated around 3 hours while the photosynthetic apparatus bursts at 8 hours to be fully accumulated at 48 hours. **C** Model for the connection of HY5 to the GLK1 genetic pathways through the action of PAP8 during the dark-to-light transition. The yellow spark represents the initial exposure of the seedling to light. Photo-initiation is mediated by PAP8 and PAP5 rapid accumulation in early photo-morphogenesis (φMG): the Pr state of phytochrome B (PHYB^Pr^) is converted in the Pfr state (PHYB^Pfr^), which enters the nucleus, accumulates in early photobodies (small green discs that regroup in late photobodies (large green discs) while promoting the destabilization of PIFs (PIF1 and 3 in particular). The red arrow in the timeline represents the albino block observed in *pap8-1* that may represent a key feature of the PAP syndrome. The dark induces ubiquitination and degradation of HY5^Ub^ impeding recognition of a *cis*-regulatory element in the promoter of PAP8 (yellow box). Light induces transcriptional activation of PAP8 in palisade cells likely through the HY5 pathway (dashed yellow arrow on PAP8 promoter); this part of the model is supported by transient assays, *in vitro* experiments, and *in vivo* genome-wide ChIP sequencing data. The cTP pre-sequence of PAP8 allows plastid import and then PAP8 assembly within the PEP-PAP complex. Using an unknown trafficking route such as travelling across the plastid envelope (?), part of the processed PAP8 pool is found in the nucleus where its action could be necessary for the PHYB-mediated transcriptional activation of GLK1 directly through HY5^30^ (dashed yellow arrow) or other light responsive factors (LR), which in turn can activate the Photosynthetic Associated Nuclear Genes (PhANGs) concomitantly to the (PEP-PAP)-driven expression of the Photosynthetic Associated Plastid Genes (PhAPGs) essential for the building of the photosynthetic apparatus (PS) in the functional chloroplast.

## Methods

### Accessions

TAIR (http://www.arabidopsis.org/) - *PAP8/pTAC6*: At1g21600; *PAP5/HMR/pTAC12*: At2g34640; *PAP10*/*TrxZ*: At3g06730; *GLK1*: At2g20570; *HY5*: At5g11260; *PHYB*: At2g18790; *EF1α*: At5g60390.

*HYH*: At3g17609; *TGA2*: At5g06950; *bZIP60*: At1g42990; *LAF1*: At4g25560; *PIL1*: At2g46970; *ELIP2*: At4g14690 *PSY*: At5g17230.

### Statistical analysis

Percentages were compared using ɛ-test whereas mean values were compared using δ-test; statistical values were confronted to the table of normal distribution (Fisher Yates: Statistical tables for biological, agricultural, and medical research (Oliver and Boyd, Edinburgh)) with α set to 0.05 or to retrieve p-values.

### Biological materials

*Arabidopsis thaliana* seeds; *pap8-1*: SALK_024431 (N524431), and Col-0: SALK_6000, were obtained from The European Arabidopsis Stock Centre NASC. *E. coli* DH5α strain (lacZ-ΔM15 Δ(lacZYA-argF) U169 recA1 endA1 hsdR17(rK-mK+) supE44 thi-1 gyrA96 relA1) was used for cloning. *Agrobacterium tumefaciens* strain C58C1 pMP90 was used for transgenesis. Rosetta™2 (DE3) (Novagen) cells were used for protein production with pBB543 (*HY5-H6*), pAG21d (PAP8^ΔcTP^-H6), pAG08 (H6-MBP-PAP5-H6) or pETM40 (MBP) see table SI1 for details.

### Plant transformation

Electro-competent *Agrobacterium* were transformed with binary plasmids containing our transgene (see table SI1) (Antibiotics: Gentamycin Rifampicin and Spectinomycin for the plasmid carrying the transgene). Strains were then used for floral dip infiltration of the significant genotypes (Medium: 2.2 g MS salts, 1 ml Gamborg’s 1000× B5 vitamins, 0.5% sucrose, 44 nM benzyl amino purine, 300 μL/L Silwet L-77). Sporophytic lethal *pap8-1* was used as the progeny of a heterozygous plant; transgenic plants were then selected to carry the mutant allele *pap8-1* (yielding albino plants in the progeny) and to carry the selection marker using the corresponding antibiotic.

### Growth conditions

Plants were grown on 1/2 MS media, sucrose and 0.8% agar. Seeds were imbibed and stratified for 2 days at 4°C, before growth at 21°C for 3 days in darkness. Afterward plants were transferred to continuous white light (30 μmol.m^−2^.s^−1^). For kinetics and organelle fractionations wild type was grown on MS medium without sucrose at 18°C. For pharmacological rescues of *pap8-1*, imbibed seeds were spread in sterile plastic boxes, containing ½ MS-media with 3% sucrose. After stratification see above, seeds where transferred to continuous white light (10 μmol.m^−2^.s^−1^) at 21°C for 7 days, before a shift to short day conditions (8 h light/ 16 h darkness) in the same light until robust rosette plants were developed. Afterwards plants were shifted to long day conditions (16 h light/ 8 h darkness) in order to induce flowering. Hypocotyl length was measured using *ImageJ* on pictures of agar plates after light treatments as indicated. For the *pap* genotypes grown in the dark or far red light, an additional 24-h growth under white light was necessary to pick the homozygotes present at a ratio of ¼. The plate was then compared to the picture to map and mark each mutant otherwise unrecognizable. True dark treatment was done after imbibition (2-3 h under white light) by a 5 min far red treatment (30 μmol.m^−2^.s^−1^) then wrapped in aluminium foil and placed in the dark at 21°C. Pigments were analysed by spectroscopy in 80% acetone. The chlorophyll content was normalized to the fresh weight corresponding to 70-110 mg seedlings and calculated using published formula (Porra, Thompson et al., 1989).

### Gene expression and protein sub-cellular localization

Transient expression in onion cells (bulb sliced to ~16 cm^2^) was conducted using the Biolistic PDS 1000/He Particle Delivery System (Biorad) (1100 psi, 10 cm traveling distance) with DNA onto 1 μm gold particles (Seashell Technology ™) following instructions. After 16 to 40 h in the dark at 24°C, the epidermis was peeled and observed by fluorescence microscopy with a Nikon *AxioScope* equipped with FITC filters and an *AxioCam MRc* camera. Pictures acquired with the Nikon’s Zen software. Confocal microscopy was performed on a Leica TCS SP2 or a Zeiss LSM800. Protein localization of stably transformed plants was examined on cotyledons or hypocotyls.

### Luciferase assay

Onion epidermal cells were transfected by micro-projectile bombardment with 0.5 μg Kar6 (p35S∷GFPer) 0.2 μg pRLC (Renilla luciferase), 1.5 μg of the luciferase reporter construct (pProm-Luc) and 1.5 μg of the trans-activating construct (p35S∷TF), kept dark 20 h, 21°C, ground in liquid nitrogen. GFP was used to restrict the transfected area for protein extraction in 1 mL of PBLuc buffer (200 mM NaPO4, pH 7, 4 mM EDTA, 2 mM DTT, 5% glycerol, 10 mg/L BSA and 1 mM PMSF) and assayed using the Dual-Luciferase® Reporter Assay System (Promega). Experiments were normalized to negative control set at 1.

### Electromobility shift assay

The DNA probe for the PAP8 promoter (oP8Box_F, GgataccaaaaatGAcGCTCttaattatttcc; oP8Box_R, ggaaataattaaGAGCgTCatttttggtatc) or the cold probe containing a canonical G-box (GbH5_F, GttctagtgtatcagaCACGTGtcgacaaactggtgg; GbH5_R, ccaccagtttgtcgaCACGTGtctgatacactagaa) was generated by annealing single-stranded oligonucleotides (with a protruding G on one 5’-end) in annealing buffer (10 mM Tris pH 7.5, 150 mM NaCl and 1 mM EDTA). 4 pmol of dsDNA was labelled by end filling with 8 pmol Cy3-dCTP and 1 unit of Klenow fragment for 1 h at 37°C, followed by enzyme inactivation at 65°C for 10 min. For each reaction, 10 nM fluorescent dsDNA was incubated with the protein in 20 μl binding buffer (10 mM HEPES pH 7.5, 1 mM spermidine, 1% glycerol, 14 mM EDTA pH 8, 0.3 mg mL^−1^ BSA, 0.25 % CHAPS, 28 ng/μL fish sperm DNA (Roche) and 3 mM TCEP). After 15 min incubation on ice, binding reactions were loaded onto native 6% polyacrylamide gels 0.5X TBE and electrophoresed at 90 V for 90 min at 4°C. Gels were scanned on a Chemidoc^XRS^ ™ imaging system (BioRad).

### Cloning

Minipreps were performed using Qiagen kits and DNA in-gel purification using GeneClean III kit (MPBio). All cloning PCR were done using Phusion™ High-Fidelity DNA Polymerase (Thermo Scientific). *PAP8* full-length open reading frame was amplified from cDNA prepared with germinating seedlings using primers as described in table SI1 and cloned in TA cloning vectors (pGem-T, Promega). Translational fusions of PAP8-GFP were obtained using *Xho*I *Bam*HI fragments inserted into pEZS-NL (Carnegie institution, Stanford): PAP8 and ΔcTP or *Xho*I-to-*Pml*I fragments for ΔNLS and ΔcTP/ΔNLS after PCR fragment cloning using oPAP8ΔNLS_PmlI. ΔNLS fragments were generated using the endogenous *Pml*I site at the 3’-end of the NLS and the PCR-based insertion of another *Pml*I site at the 5’-end; then NLS was clipped off using *Pml*I and backbone ligation. NLSm5 was generated using PCR-based site-directed mutagenesis (see table S1 for primers). For plant transgenesis, ORFs were cloned into pBB304e (pPAP8∷GUS^10^) or derivatives of pART27(Blanvillain, Wei et al., 2011).

### Mutant characterization and RT-PCR

gDNA preparation: leaf tissues were ground in 1.5-mL reaction tubes, then homogenized in 400 μL of EB buffer (200 mM Tris HCl pH 7.5, 250 mM NaCl, 25 mM EDTA, 0.5% SDS). After 5 minutes at 10 000 *g*, 400 μL of supernatant was added to 400 μL isopropanol. After 10 minutes at 10 000 *g*, the pellet was washed with 750 μL of EtOH 80% and then dried. DNA was then suspended in 50 μL of water. The PCR was done with indicated primers. For RT-PCR, the RNeasy plant minikit (Qiagen) was used; RNA samples were treated with RNAse free DNAase. The RTs were performed using 2 μg of RNA, SuperScript IV VILO kit (invitrogen), dT_18_ primer, 1^st^ strand buffer and RNase inhibitor. The RT programme was set at 65°C for 5 min, 5°C for 1 min, then after addition of RT mix, 42°C for 50 min and 70°C for 10 min. The PCR was done with indicated primers on 0.5 μL of cDNA. Absence of genomic DNA was checked by PCR on EF1α. oPAP8_rtp_F, tggtggtgatggagatatcg; oPAP8_rtp_R, tttgagacactgaagtctcg; op8i2_R, aaggaagtctcagaacaacgc; oLBb1.3, attttgccgatttcggaac; oE3_R, tagtcactcattgcacatcg; EF1α: F, caggctgattgtgctgttcttatcat; R, cttgtagacatcctgaagtggaaga. GLK1: Frtpcr, cacatgaacgcttcttcaacg; Rrtpcr, tgtagctctggtgtccaatcc. qPCR on GLK1 and GLK2 was performed using Power SYBR Green Master Mix (ThermoFisher Scientific). Primer sequences were designed with Quantprime. Their efficiencies were between 90 and 110%, and they did not amplify genomic DNA. oEF1α_qF, tgagcacgctcttcttgctttca; oEF1α_qR, tgtaacaagatggatgccaccacc; oGLK1_qF, ttctaccgccatgcctaatccg; oGLK1_qR, actggcggtgctctaaatctcg; oGLK2_qF, agcatcggtgttcccacaagac; oGLK2_qR, tcgagggatgaatgtcgatggg.

### Protein production. HY5

Rosetta2 cells were grown overnight in 50 mL LB with 100 μg/mL of carbenicillin and 34 μg/mL of chloramphenicol at 37°C. 1 L of LB + antibiotics was then inoculated and cultivated at 37°C to 0.1 OD_600_. At 0.6 OD_600_, the temperature was decreased to 16°C and 0.5 mM of isopropyl β-D-1-thiogalactopyranoside was added. After an overnight induction, cells were harvested by centrifugation at 5,500 *g*, for 25 min, at 4°C. The cell pellet was resuspended in 30 mL of lysis buffer: 50 mM Tris HCl, pH 8.0, 0.5 M NaCl, 20 mM imidazol pH 8.0 with a Complete™ Protease inhibitor Cocktail tablet (Roche). The lysate was centrifuged at 15,000 *g*, for 40 min, at 4°C. The purification was performed at 20°C. After filtration, the supernatant was applied onto a NiNTA column in 50 mM Tris HCl, pH 8, 0.5 M NaCl, 20 mM imidazol pH 8. HY5 was eluted in 50 mM Tris HCl, pH 8, 0.5 M NaCl and 300 mM imidazol. After dialysis in 50 mM HEPES pH 7, 0.5 M NaCl, HY5 was concentrated using an Amicon Ultra 15 mL centrifugal filter and a 10-kDa-membrane cut-off before loading on a Superdex 75 10/30 and eluted with 25 mM HEPES pH 7, 50 mM NaCl. **PAP8.** Production as above except that 10 mM β-mercaptoethanol was added to lysis and elution buffers; 1 mM DTT added to dialysis buffer. PAP8 was loaded on a Superdex 200 10/30 and eluted with 10 mM Tris HCl pH 8, 50 mM NaCl, 5 mM DTT. Rabbit polyclonal antibodies against PAP8 were produced by ProteoGenix. In Western blots PAP8 is detected at ~38 kDa; which is 7 kDa larger than the theoretical MW of processed PAP8 (31.1 kDa) without its transit peptide (6.2 kDa). With the MW correction on negative charges (D/E) using the linear correlation of Guan *et al*., (Guan *et al*., 2015) (equation y = 276.5x - 31.33, where x is the ratio of acidic AA (D+E) and y the average of delta MW in Da per AA), the high occurrence of D+E in PAP8 (58/269=0.216) then causes a calculated retardation of 7.6 KDa, which explains the apparent MW of PAP8 observed in western blot. ^15^**N-PAP8** was produced in minimum medium M9 containing 1g/L ^15^NH_4_Cl (M9-^15^N). 5 ml of LB + antibiotics were inoculated with cells containing pAG21d. After 10h of growth, 1 mL was added to 100 ml of M9-^15^N + antibiotics. At 2 OD_600_ (16h), the culture was centrifuged at 4000 *g* and the pellet was used to inoculate 1L of M9 + antibiotics. Culture, induction and purification were done as described for PAP8. **MBP-PAP5**. Culture and overexpression was done as for PAP8; 100 μg/mL of kanamycin was used instead of carbenicillin. 20 mM β-mercaptoethanol was added to lysis buffer. The lysate was centrifuged at 15,000g, for 40 min, at 4°C. The purification was performed at 20°C. The supernatant was applied onto an amylose column in 20 mM Tris HCl pH 7.5, 200 mM NaCl, 1 mM EDTA, 10 mM β-mercaptoethanol. MBP-PAP5 was eluted in 20 mM Tris HCl pH 7.5, 200 mM NaCl, 1 mM EDTA, 10 mM β-mercaptoethanol, and 10 mM maltose. MBP-PAP5 was then concentrated with an Amicon Ultra 15 mL centrifugal filter and a 30-kDa-membrane cut-off before loading on a HiLoad 16/60 Superdex 200 and eluted with 10 mM Tris HCl pH 8.0, 50 mM NaCl, 5 mM DTT. **MBP.** Culture and overexpression in LB followed the same procedure than for PAP5. The cell pellet was resuspended in 30 mL lysis buffer. The lysate was centrifuged at 15,000g, for 40 min, at 4°C. The purification was performed at 20°C. The supernatant was applied onto an amylose column in 20 mM Tris HCl pH 7.5, 200 mM NaCl, 1 mM EDTA. MBP was eluted in 20 mM Tris HCl pH 7.5, 200 mM NaCl, 1 mM EDTA, and 10 mM maltose. MBP was then concentrated with an Amicon Ultra 15 mL centrifugal filter and a 10-kDa-membrane cut-off before loading on a HiLoad 16/60 Superdex 75 and eluted with 10 mM Tris HCl pH 8.0, 150 mM NaCl, 10% glycerol. The pools containing pure HY5, PAP8, PAP5 or MBP were concentrated using Amicon centrifugal filter.

### Protein extraction and Western immuno-detection

5-day-old *Arabidopsis* (50 mg, approx. 100 seedlings) are collected and homogenized using a Precellys™ tissue homogenizer (3 × 20 s, 9,300 *g*, break 30 s) in 100 μL of denaturing extraction buffer (DEB: Tris HCl 100 mM pH 6.8, Urea 8 M, EDTA/EGTA 10 mM, DTT 10 mM, protease inhibitor (Roche) 1 tablet/10 mL, 100 μL glass beads (diameter 4-6 mm). The samples are centrifuged (10 min, 4°C, 9,300 *g*). The total soluble protein samples (TSP) are titrated by Bradford assay before mixing in Laemmli buffer (Tris HCl 100 mM pH6.8, Glycerol 10%, SDS 2%, DTT 50 mM, Bromophenol Blue 0.25%) and heated 10 min at 80°C. TSP were separated by SDS-PAGE and transferred on nylon membrane (Biorad). The membrane was block in TBS, Tween 0.1%, non fat dry milk 5% w/v. The membrane was probed in TBS Tween 0.1%, with different primary antibodies against PAP8 (this study), PAP5 (PhytoAB, Ref. PHY0389), Histone H3 (Agrisera, Ref. AS10710), RbcL (Agrisera, Ref. AS03037), HY5 (PhytoAB, Ref. PHY0264), PHYB (PhytoAB, Ref. PHY0750), PIF1 (PhytoAB, Ref. PHY0830), PIF3 (PhytoAB, Ref. PHY0063); dilutions H3, RbcL: 1/10,000; others: 1/5,000. Membranes were washed (5 times, 5 min in a TBS-Tween 0.1%); secondary antibody, Goat anti-Rabbit conjugated with a Horse Radish Peroxidase was used at a dilution of 1/5,000. Signal was detected using a chemiluminescent substrate (Biorad, ECL kit).

### Organelles fractionation

5-day-old *Arabidopsis* seedlings, exposed to light or dark, were homogenized in liquid N_2_. The powder was dissolved in a cold native extraction buffer (NEB: Tris HCl 100 mM pH 7.4, glycerol 25%, KCl 20 mM, EDTA 2 mM, MgCl_2_ 2.5 mM, Sucrose 250 mM, DTT 5 mM, protease inhibitor Roche™, 1 tablet/ 50 mL) at a ratio of 1:3 (w/v). The extract was filtered through 3 layers of miracloth and one layer of nylon (100 μm) centrifuged (10 min at 1500 *g*, 4°C). The supernatant was deposited on percoll 80% and centrifuged (swinging rotor, 5 min at 2,300 *g*, 4°C) to remove the pellet of starch. The supernatant was loaded on 35% percoll and centrifuged (swinging rotor, 5 min at 2,300 *g*, 4°C) to separate swimming plastids from the pellet of nuclei. The nuclei were washed two times with a plastid-lysis buffer (NEB + 2% triton) followed with centrifugation (5 min at 1500 *g*, 4°C). Fractions, corresponding to plastids or nuclei, were suspended in DEB shaken on a vortex (10 min at 4°C) before centrifugation (10 min at 9,300 *g*, 4°C). The TSP samples were subjected to western blot analysis as above.

### NMR spectroscopy

The ^1^H-^15^N spectrum of the ^15^N-labelled PAP8 alone was recorded at 300 K using a BEST-TROSY experiment on a protein sample concentrated at 100 μM in 10 mM Tris buffer containing 50 mM NaCl and 5 mM DTT in a 95:5% H_2_O:D_2_O solvent at pH 8.0 for 49 minutes. The same experiment was performed on ^15^N-labelled PAP8 at a concentration of 88 μM in presence of an unlabelled PAP5-MPB construct in stoichiometric conditions increasing the number of scans to reach an experimental time of 13 hours. The control experiment involving ^15^N-PAP8 and unlabelled MPB also in stoichiometric conditions was performed to check the interaction assumption between the 2 proteins. The following NMR experiments ^1^H-^15^N BEST-TROSY(Favier, Brutscher et al., 2011), TRACT (to estimate the global correlation time) (Lee, Hilty et al., 2006) and DOSY experiments (for measuring the translational diffusion) (Morris & Johnson, 1992) were recorded on a Bruker *AVANCE™* III spectrometer operating at ^1^H-frequency of 700 MHz and equipped with a triple resonance pulsed field gradient cryo-probe.

## Acknowledgments

We acknowledge the platforms of the Grenoble Instruct-ERIC centre (ISBG; UMS 3518 CNRS-CEA-UGA-EMBL) within the Grenoble Partnership for Structural Biology (PSB). Platform access was supported by FRISBI (ANR-10-INBS-05-02) and GRAL, a project of the University Grenoble Alpes graduate school (Ecoles Universitaires de Recherche) CBH-EUR-GS (ANR-17-EURE-0003). IBS acknowledges integration into the Interdisciplinary Research Institute of Grenoble (IRIG, CEA). The work was supported by the *Agence National de la Recherche* (grant PepRegulChloro3D), the *Deutsche Forschungsgemeinschaft* to T.P. (PF323-5) and the AGIR programme of *Université Grenoble-Alpes* (UGA) to R.B. The project received further support by institutional grants to the *Laboratoire de Physiologie Cellulaire et Végétale* by Labex Grenoble Alliance of Integrated Structural Biology (GRAL) and ANR-17-EURE-0003. We thank F Barneche for the ChIP-seq analysis at the PAP8 locus; E Monte and G Toledo-Ortiz for *pifq* seeds; A Guerrero-Criado, R Toutain, S Coveley for their help at the bench. We thank E Thevenon in the Parcy lab for advice on fluorescent labelling EMSA. We thank S Lerbs-Mache for critical reading. We express our gratitude in memory of D Grunwald for his help on confocal imaging.

## Author contributions

TP, DC, and RB designed research. ML, FXG, AI, KP, LC, MC, RR, FC, DC, RB, performed research. DC, TP, RB analysed data. EBE contributed mass spectrometry data. AF, PG contributed NMR data. TP and RB wrote the paper with contributions from ML, FXG, DC, AF, PG and EBE. All authors approved the paper.

The authors declare that they have no conflict of interest.

## Extended materials

### Figure Legends

**Fig. S1| (Related to Fig. 1) Genetic analysis of the *pap8-1* mutant. A** The two phenotypic classes [WT] and [albino] from a *pap8-1*/+ segregating progeny and their ratio in **B** calculated from N=604 plants. **C** Box plot (Min, 1^st^ quartile, median, 3^rd^ quartile, Max) of cotyledon width 5 days after germination (DAG) in wild type and albino plants grown *in vitro* under 30 μmol.m^−2^.s^−1^ of white light. **D** Functional complementation of *pap8-1*: phenotypes of *pap8-1* homozygous plant grown *in vitro*, and two representative plants of wild type or *pap8-1/*pP8∷PAP8 (line L35 or line L49) grown on soil. **E** Spectrophotometric analysis of pigments: absorption spectra of acetone-soluble extracts from seedling grown *in vitro* 3 days in the dark (D) or 3 days in the dark plus 30 hours of white light (L) Col-0, wild type; p8/p8, homozygous mutant *pap8-1*; L35 and L49, two lines of *pap8-1/*pP8∷PAP8; n.a., not applicable. Absorbance was normalized to fresh weight (FW); Chla, chlorophyll a; Chlb, chlorophyll b; Car, carotenoids.

**Fig. S2| (Related to Fig. 1 and 3) Flow chart of the *pap8-1* functional complementation test.** *p8-1*, *pap8-1* allele associated with *nptII*, *neomycin phosphotransferase II* marker used to select plants resistant to kanamycin (Kan^R^) although this resistance is partially lost in *pap8-1*. **A** Heterozygosity test on the progeny of one plant, if 1/4 of albino plants appear then the tested seeds were used for floral dip with an allelic frequency of 1/3 for *pap8-1*, *Agrobacterium*-mediated transformation; *hptII, hygromycin phosphotransferase II* gene to select resistant plants (Hyg^R^) from those that are untransformed and sensitive (Hyg^S^). Selection under low red light (660 nm at 8 μmol.m^−2^.s^−1^) allowing for rapid elongation of Hyg^R^ plants. The etiolating response cannot be used with PBG that causes a strong de-etiolated phenotype. Primary transformants (T1) were PCR-selected for the presence of the tested *goi* (gene of interest), presence of the *pap8-1* allele, and the wild-type allele. The yellow scenario represents a successful complementation test in T1 (FC); the albino plant in brackets may be retrieved in some tests at a low ratio (1/6 of all T1) as negative for the complementation test (noC). The white box represents a common event of interest (1/3 of all T1 carrying the *goi* and one allele *pap8-1*). Conclusion is made after testing the genetics in the T2 generation. **B, C** Heterozygosity test to retrieve doubly homozygous (**C**) complemented T3 line with the pPAP8∷PAP8 transgene (p8P8).

**Fig. S3| (Related to Fig. 2) PAP8 promoter analysis. A** Cartoon representing the different annotations of the promoter and the plot of local % of W= (A or T) in steps of 10 nucleotides of the nucleic acid sequence. Red, 5’-untranslated regions; pink, coding sequence of At1g21610 upstream gene on the reverse strand; **B** Table of the occurrences (occ.) of binding sites detected for the different transcription factors (TF) families using PlantPan3^26^. The promoter was broken down to two regions (−497 to −97) and (−97 to +1); **C** Selected binding sites represented on the (−497 to +63) PAP8 promoter sequence, >, plus strand; <, minus strand; <-->, element detected on both strands; the yellow box depicts the near palindromic bZIP element found at −97; annotations for individual elements are given according to Table S2.

**Fig. S4| (Related to Fig. 2) Transactivation test in onion epidermal cells using the dual luciferase reporter assay.** TF, transcription factor; Luc, luciferase; glow is the emission of photons after the protein samples have been supplemented with the luciferase-type specific substrate. Activity of the tested transcription factor calculated as the relative value of Firefly glow / *Renilla* glow to that of the reference sample.

**Fig. S5| (Related to Fig. 3) A Immunoblots showing the levels of PAP8 and PAP5** in Col0, *pap8-1* (*p8-1*) or *pap5-2* (*p5-2*) respectively, using the recombinant proteins PAP8 (rP8) and PAP5 (rP5) as controls. Proteins on PAGE were detected by instant blue to display equal loading. **B** PAP8 immunodetection in fractions obtained during organelles enrichment; rP8, as above; S1, soluble fraction in the supernatant after centrifugation of the blender-disrupted cellular sample; Tot, total plant extract; C+N, pellet containing organelles, mostly chloroplasts and nuclei separated from S1; H3 antibody used to evaluate nuclear enrichment and a Coomassie staining presented as loading control.

**Fig. S6| (Related to Fig. 3) Sub-cellular localization of PAP8 variants. A** Transient assay in onion epidermal cells. FL, full-length ORF; -G, translational fusion with GFP; ΔNLS, deletion of the NLS; ΔcTP, deletion of the cTP. Onion cell co-transfected with the corresponding variant fused to GFP and a plastid control fused to RFP (PAP10, PAP10-RFP or RecA, RecA-RFP) Merge, merged channels; DIC, differential interference contrast microscopy pictures to reveal the position of the nucleus within the cell when fluorescent nuclei were observed. **B, C** Confocal imaging of stably expressed CaMV35S∷PAP8^ΔcTP^-GFP (b) or pPAP8∷PAP8^Δnls^-GFP (**C**) in cotyledons of *Arabidopsis thaliana*; white arrowheads indicate nuclei; green arrowheads indicate sub-plastidial localization. Observations similar to (**C**) were recorded for pPAP8∷PAP8^FL^-GFP. **D** Confocal imaging of stably expressed CaMV35S∷PAP10-GFP in cotyledons of *Arabidopsis thaliana* showing the PEP-PAP complex; white arrowheads indicate nuclei; **E-H** PAP8-GFP in stromules of onion epidermal cells expressed alone (**E**) or in co-localization with PAP10-GFP (**F, G, H**); stromules (str) are only marked by PAP8-GFP.

**Fig. S7| (Related to Fig. 3) PAP8 and its variants fused to GFP. A** Schematic illustration of the GFP-fused PAP8 variants used for sub-cellular localisation and functional complementation tests FL-G, *PAP8* full-length ORF translationally fused to GFP; Δnls-G, deletion of the NLS, ORF fused to GFP; ΔcTP-G, deletion of the cTP, ORF fused to GFP; ΔΔ-G combined deletion of NLS and cTP in the ORF fused to GFP. The variants are expressed under the control of the CaMV35S promoter or under the native PAP8-1.1-kb promoter. Plasmid identification and relevant restriction sites for cloning are given in light grey (see table S1). **B** Transgenic lines obtained with the constructions described above. Three phenotypic classes have been recorded corresponding to albino (white squares) pale green (light green squares) and green plants (green squares). Col-0, wild type; *pap8-1/+*, heterozygous mixture; n, total number of recorded plants. **C** Functional complementation output. Hygromycin resistant plants were transferred on soil and grown under long day conditions (16 h light / 8 h dark; ~70 μmol.m^−2^.s^−1^) at 21°C and 60 % humidity. Genomic DNA was isolated from true leaves and used for genotyping. The presence of the *pap8-1* allele was confirmed using the primer ortpF/oLBb1.3. PAP8 wild-type allele tested with ortpF/op8i2_R. The insertion of the transgene of interest was tested with ortpF/oE3R. The number of the tested plants (Hyg^R^#T1), the number of double heterozygous plants (p8-1/+;TG/−) and the number of sesqui-mutant plants (*p8-1/p8-1*; TG/−) are depicted for each construction; p8P8 presented as positive control. None of the 39 T1 plants with pP8∷PAP8-GFP, were photosynthetic and homozygous *pap8-1*. Therefore, two doubly heterozygous (*pap8-1*/+; TG/−) expressing GFP were tested for their segregation pattern (Line 1: 34% albino, n=169 and Line 2: 24% albino, n=199) and compared to that of *pap8-1*/+ (28% albinos, n=99). In absence of statistical difference between the samples (ɛ = 0.102<<1.96 for α=0.05; ɛ-test, Fisher Yates), PAP8-GFP was declared not functional as opposed to PAP8.

**Fig. S8| (Related to Fig. 4) Hemi-complementation test in *pap8-1*.** Phenotype of *pap8-1* transformed with pPAP8∷PAP8^NLSm5^. WT, 5-week-old Col-0 control; [DG], Delayed Greening phenotype observed for the partial rescue of *pap8-1* mutant expressing PAP8^NLSm5^ under its endogenous promoter; pictures depict three 15-week-old plants in which the alteration of the greening corresponds to the emergence of white leaves that slowly acquire the photosynthetic apparatus; plants #7, #31 and #57 are siblings of the same genotype (Ai10#34). Bars equal 10 mm.

**Fig. S9| A** Hypocotyl length measurements of genotypes grown under different light sources. Dark, true dark treatment; FR far red light (low fluence approx. 10 μmol.m^−2^.s^−1^; peak at 730 nm +/− 10 nm); R, red light (8 μmol.m^−2^.s^−1^ peak at 660 nm +/− 10 nm); L, white light 30 μmol.m^−2^.s^−1^. Whereas *pap5-2* and *pap8-1* are statistically undistinguishable (δ = 0.791 << U_α=0,05_ = 1.96; Fisher Yates) both *pap5-2* and *pap8-1* show significant hypocotyl differences compared to wild type (δ = 5.374; δ = 5.061 respectively so that both p-values < 10^−6^). **B** Images of representative seedlings.

**Fig. S10| (Related to Fig. 6) Bimolecular fluorescence complementation test.** Using PAP8^ΔcTP^-NY (P8Δc-NY) and PAP5^ΔcTP^-YC (P5Δc-YC) and the reverse combination; PAP8 tested positive using P8Δc-NY with P8Δc-YC.; arrowheads indicate nuclei. P8Δc-NY, P5Δc-NY, P8Δc-YC and P5Δc-YC tested negative using respectively both split YFP fragments alone (ϕ-YC or ϕ-NY). PAP10-RFP (P10-RFP) used as internal control for transfection efficacy knowing YFP signal could be absent. The number followed with a letter refers to the pictured cell and the ratio depicts the number of green-fluorescent cells over red-fluorescent cells. White arrowheads indicate the nuclei and the scale bar equals 20 μm.

**Fig. S11| (Related to Fig. 6) Variations of the diffusion coefficient and the peak intensities of PAP8 in function of the PAP5/PAP8 concentration ratios.** Triangles, normalized values of the translational diffusion coefficient of PAP8. Squares, normalized intensities of peaks in the 7.5–8 ppm range (structured). Circles, normalized intensities of peaks outside the 7.5–8 ppm range (unstructured).

**Table S1| Plasmid overview and the primers used for their cloning.** bR: bacterial resistance to antibiotics; carb, carbenicillin; kan, kanamycin; spec, spectinomycin. PlantR, plant resistance used as the selection marker and brought by the binary vector: Hygro, hygromycin. Genotype used for this study: Col-0 or wt, wild type Columbia; *p8-1*, heterogeneous progeny from individual heterozygotes *pap8-1/+* used for transformation. Primers are given from 5’ to 3’. n.a. not applicable. fc1 and fc2, each promoter driving PAP8 coding sequence (TG) was tested in functional complementation: two lines doubly heterozygous *pap8-1*/+; TG/− segregated around 1/16-albino plants. (fc1) pP8^−257^∷PAP8, Line 1 (n=95, ɛ=0.76<<1.96); Line 2 (n=62, ɛ=0.06<<1.96). (fc2) pP8^−97^∷PAP8, Line 1: n=389, ɛ=0.34<<1.96; Line 2: n=392, ɛ=0.32<<1.96 (ɛ-test, α=0.05; Fisher Yates).

**Table S2| Output from PlantPan3: Search for transcription factors binding sites.** Annotations as on the corresponding website; position is given relative to the transcriptional initiation site.

